# Cell volume tunes macrophage innate inflammatory responses through promoting type I interferon signalling

**DOI:** 10.1101/2024.07.15.603505

**Authors:** James R Cook, Tara A. Gleeson, Stuart M. Allan, Catherine B. Lawrence, David Brough, Jack P. Green

## Abstract

Macrophages are key effectors in co-ordinating inflammatory and immune responses to threats to the host. How macrophages decipher diverse danger signals to tailor inflammatory responses remains an unanswered question. Cell volume control is critical for normal cellular function. Disturbances in extracellular and intracellular homeostasis induce changes in cell volume, but the impact of disruptions in cell volume in controlling macrophage inflammatory responses is poorly understood. Here, we discover that macrophages use cell volume control as a bona fide danger sensing mechanism to promote and augment inflammatory responses. Using macrophages deficient in the volume regulated anion channel (VRAC), which lack cell volume control under hypo-osmotic conditions, we show that disruptions in cell volume are sensed by macrophages to drive a large transcriptomic response and induction of inflammation. Cell volume disruption, particularly loss of cell volume control, induces type I interferon signalling through a DNA– and STING-dependent mechanism, but independent of cGAS and 2’3’cGAMP transport. Further, we found that cell volume changes synergise with diverse pathogen-mediated signalling to augment type I interferon responses and exacerbate the cytokine storm in a mouse model of hyperinflammation. Our findings highlight cell volume as an important regulator in shaping inflammatory responses, adding to our understanding of how macrophages sense complex danger signals and threats.

## Introduction

The innate immune system is an early line of defence against invading pathogens and tissue damage and functions to promote pathogen clearance and wound healing. Macrophages represent the cellular frontline of the innate immune system and are specialised to detect danger signals, features of infection (pathogen associated molecular patterns; PAMPs), cellular damage (damage associated molecular patterns; DAMPs) or disturbances in extracellular and intracellular homeostasis to initiate and sustain inflammatory responses^1^. The repertoire of danger signals sensed by macrophages is diverse and extensive highlighting a crucial role for macrophages in danger sensing^2,3^. Furthermore, the nature of the inflammatory response initiated varies depending on the type of danger signal sensed to resolve the threat most appropriately. The mechanisms utilised by immune cells to decipher the many different danger signals in the inflammatory environment to promote an optimal response is an area with many unanswered questions.

A relatively unexplored contributor to the tuning of inflammatory responses is the regulation of macrophage cell volume. Cell volume control is critical for normal cellular function, where excessive changes in cell volume threaten cell survival and homeostasis. Cell volume is constantly regulated by proliferative and metabolic processes where loss of cell volume control, and resulting cell volume dysfunction, is commonly associated with features of the inflammatory environment, such as oedema^4^, hypoxia and metabolic dysfunction^5^, with cell swelling a common feature of pro-inflammatory cell death^6^. Cells use mechanisms to regulate cell volume, including ion channels such as the volume regulated anion channel (VRAC) in vertebrates. In swollen cells, VRAC drives chloride ion (Cl^-^) efflux to promote water efflux to restore cell volume, through a process known as regulatory volume decrease (RVD)^7,8^. Loss of VRAC results in dysregulated cell swelling following hypotonic shock through an inability to undergo RVD^7,9^. Since the discovery of LRRC8 family proteins as the sub-units of VRAC^7,8^, it is now known that as well as cell volume regulation, VRAC is implicated in diverse cellular functions such as cell migration, proliferation, apoptosis, cell-cell communication, and in immunity^10^. VRAC and the RVD are emerging as important regulators of inflammatory signalling in immune cells by regulating activation of the NLRP3 inflammasome^9,11^, and cGAS-STING pathways^12–14^. Furthermore, treatment with hypertonic solutions, which prevents cell swelling, is effective at alleviating inflammation *in vivo*, including in LPS-induced sepsis^15^, intracerebral haemorrhage^16^, pulmonary injury due to ischaemia-reperfusion^17^, and acute respiratory distress syndrome (ARDS)^18^. However, despite the reported indications that cell volume and VRAC are involved in inflammatory signalling, the basic biological mechanisms of how the regulation of cell volume shapes inflammation are currently unknown.

Here, we demonstrate that regulation of macrophage cell volume shapes the innate immune response. By using conditional VRAC knockout (KO) macrophages, we revealed that loss of cell volume control resulted in an altered transcriptional response to hypotonicity, including an induction of inflammatory signalling promoting an antiviral cell state. We show that swelling of macrophages drove type I interferon signalling through a STING-dependent pathway identifying a new mechanism of inflammation following cell swelling. We show that VRAC-deficient macrophages exhibited altered cytokine responses following stimulation of PAMPs in hypo-osmotic conditions, promoting enhanced type I IFN responses, suggesting that regulation of cell volume influences the nature of the immune response. Furthermore, we provide evidence that the regulation of macrophage cell volume contributes to inflammation *in vivo* by observing exacerbated inflammatory responses in conditional CX3CR1 cell specific VRAC knockout mice in a model of CpG-DNA-induced hyperinflammation. Therefore, we propose that the ability of a macrophage cell to regulate its volume is critical to shaping subsequent inflammatory responses.

## Results

### Altered regulation of macrophage cell volume initiates transcriptomic reprogramming and induction of inflammation

We hypothesised that perturbations in the ability of a cell to control its cell volume would act as a signal to promote inflammation. We have previously described and characterised macrophages from mice lacking LRRC8A (a subunit essential to all VRAC channels^7,8^) in CX3CR1^+ve^ cells (*Cx3cr1*^Cre^ *Lrrc8a*^fl/fl^ mice, termed throughout as VRAC KO macrophages), which lack the ability to undergo RVD in response to hypotonic shock, and as a result stay swollen^9,19^. Therefore, VRAC KO macrophages are an ideal tool to investigate how a loss of cell volume control regulates inflammatory signalling. We have previously shown that upon exposure to severe hypotonic conditions (117 mOsm kg^−1^), that the macrophage RVD response induces VRAC-dependent activation of the NLRP3 inflammasome^9^, providing evidence that control of cell volume influences inflammatory signalling. However, these are extreme hypotonic conditions, leading us to question how macrophages would respond to less severe changes in cell volume in circumstances where the inflammasome is not activated. To this end, we first performed bulk RNA-sequencing on WT and VRAC KO macrophages following less severe cell swelling to assess what, if any, transcriptomic changes occurred. Primary bone marrow-derived macrophages (BMDMs) generated from WT and VRAC KO littermate mice were treated with hypo-osmotic media (50% v/v H_2_O in DMEM, ∼170 mOsm kg^−1^) to induce cell volume changes or left untreated (Fig 1A). Cre control (*Cx3cr1*^cre^ *Lrrc8a*^WT/WT^) macrophages were also included in the untreated group (Fig 1A). In parallel, we confirmed that the VRAC KO macrophages cultures used for RNA-sequencing had lost the ability to undergo RVD in response to hypo-osmotic media (170 mOsm kg^−1^) (Supplementary Fig 1A,B). Principal component analysis (PCA) of the gene-wise read counts showed close clustering of unstimulated macrophages from WT, macrophage VRAC KO and Cre control mice, indicating that these cells exhibited very little differences under resting conditions (Fig 1B). By contrast, WT and VRAC KO BMDMs treated with hypo-osmotic media separated from the untreated groups along PC1, indicating that treatment with hypo-osmotic media had induced transcriptomic changes (Fig 1B). Interestingly, VRAC KO macrophages stimulated with hypo-osmotic media exhibited even greater separation from WT cells, suggesting that the inability to undergo RVD had further altered the transcriptomic response. We then performed differential expression analysis to identify which genes were altered between groups. Matching the PCA analysis, there were essentially no differentially expressed genes (DEGs) between genotypes under resting conditions (WT vs VRAC KO: 2 genes, WT vs Cre control: 4 genes, VRAC KO vs Cre control: 6 genes). However, there was a large transcriptomic response following hypo-osmotic treatment, with 6315 DEGs (3473 upregulated, 2842 downregulated) between unstimulated and hypo-osmotic media treated WT macrophages (Fig 1C). Similarly, VRAC KO macrophages had 7171 DEGs (3748 upregulated, 3423 downregulated) between unstimulated and hypo-osmotic treated cells (Fig 1D). Comparing between hypo-osmotic treated WT and VRAC KO BMDMs revealed 3563 DEGs (1166 upregulated, 2397 downregulated) (Fig 1E). These data suggest that regulation of cell volume elicits large changes in patterns of gene transcription and that many of these changes are regulated by VRAC.

**Figure 1.**
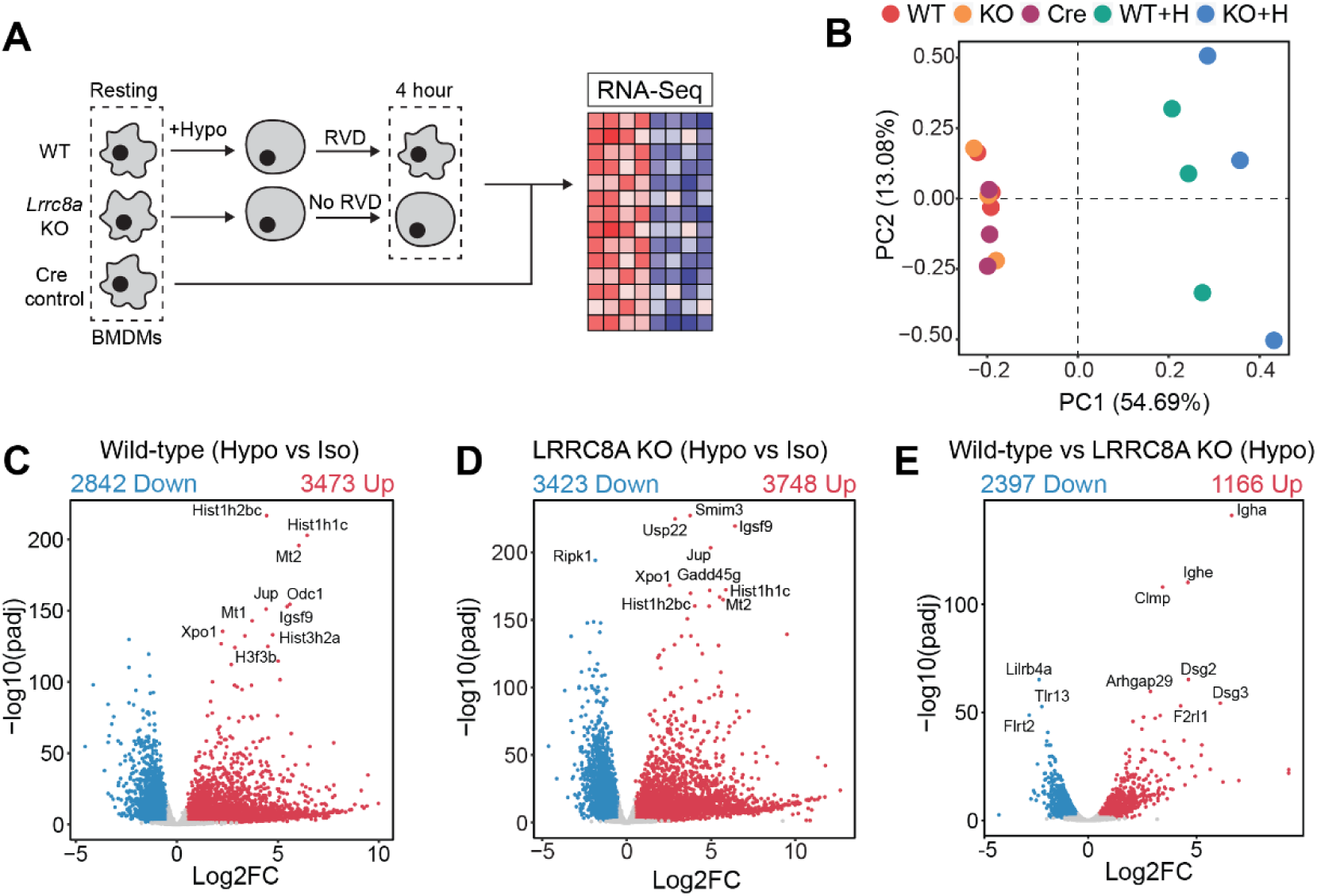
– Changes in macrophage cell volume induce a large transcriptomic response that alters diverse cellular processes. (**A**) RNA-sequencing was performed on resting bone marrow derived macrophages (BMDMs) from wild-type (WT), VRAC knockout (KO) or CX3CR1 Cre expressing controls (Cre), and WT and KO BMDMs incubated in hypo-osmotic media (50% v/v H_2_O in DMEM) for 4 hours. **(B)** Principal component analysis of the gene-wise read counts from (A). **(C-E)** Volcano plots of differentially regulated genes (DEGs) in: resting WT macrophages vs WT in hypo-osmotic media (C), resting KO macrophages vs KO in hypo-osmotic media (D), WT macrophages in hypo-osmotic media vs KO macrophages in hypo-osmotic media (E).

### Cell swelling induces an antiviral response through STING-dependent type I IFN signalling

Since we hypothesised that perturbations in cell volume would drive inflammatory responses in macrophages, we examined the expression of cytokines within the hypo-osmotic treatment RNA-seq dataset in more detail. Hypo-osmotic cell swelling induced a widespread induction of multiple cytokines, including IL-1α, IL-1β, IL-6, IL-33, TNF, CCL2 and CD40 amongst others (Fig 2A). We observed that hypo-osmotic cell swelling caused an induction of the type I IFNs, particularly IFNβ, but not type II IFN (IFNγ), or type III IFNs, (IFNλ) (Fig 2B). Further, whilst type I IFN expression was mostly absent at baseline in WT and VRAC KO BMDMs, VRAC KO BMDMs exhibited a larger induction of type I IFNs than WT BMDMs following hypo-osmotic treatment (Fig 2B). Treatment with hypo-osmotic media also induced expression of interferon stimulated genes (ISGs), which are commonly transcribed downstream of IFN signalling^20^, although there was no difference in ISG upregulation between WT and VRAC KO macrophages following 4 hours of incubation in hypo-osmotic media (Fig 2C). We then examined the kinetics of type I IFN signalling in WT and VRAC KO BMDMs using qRT-PCR for *Ifnb* and the ISG *Cxcl10*. Whilst there was no noticeable up-regulation of *Ifnb* in WT BMDMs following treatment with hypo-osmotic media, a small upregulation of the ISG *Cxcl10* was present at 4 hours and enhanced further at 6 hours (Fig 2D,E). Interestingly in VRAC KO BMDMs, *Ifnb* levels were up-regulated following 4 hours of hypo-osmotic treatment, which was further augmented at 6 hours (Fig 2D). *Cxcl10* was also upregulated from 4 hours in hypo-osmotic media treated VRAC KO BMDMs, which was significantly higher than that of WT BMDMs at 6 hours (Fig 2E). The expression of the ISGs *Rsad2* and *Ccl7,* and the inflammatory cytokine *Il1b*, were also up-regulated at 6 hours following hypotonicity-induced cell swelling in WT and VRAC KO BMDMs, with a significantly enhanced response seen in VRAC KO BMDMs (Fig 2F,H). Together, these data suggest that enhanced IFNβ signalling at earlier time-points could be promoting greater ISG expression at later time-points. Whilst upregulated mRNA expression of *Ifnb* occurred following treatment with hypo-osmotic media, IFNβ was not detected extracellularly, potentially due to the low sensitivity of the ELISA, suggesting hypo-osmotic media induced only a mild interferon response compared to stimuli such as nucleic acid addition or viral infection (data not shown). However, the induction of viperin (translated from *Rsad2)* was detectable from 4 hours and was slightly, but significantly, enhanced in VRAC KO BMDMs at 8 hours (Fig 2I,J) matching mRNA data at 6 hours (Fig 2F). These data demonstrate that changes in the ability of the macrophage to control its cell volume cause inflammation, particularly type I IFN signalling, which is enhanced when cells are unable to undergo RVD.

**Figure 2.**
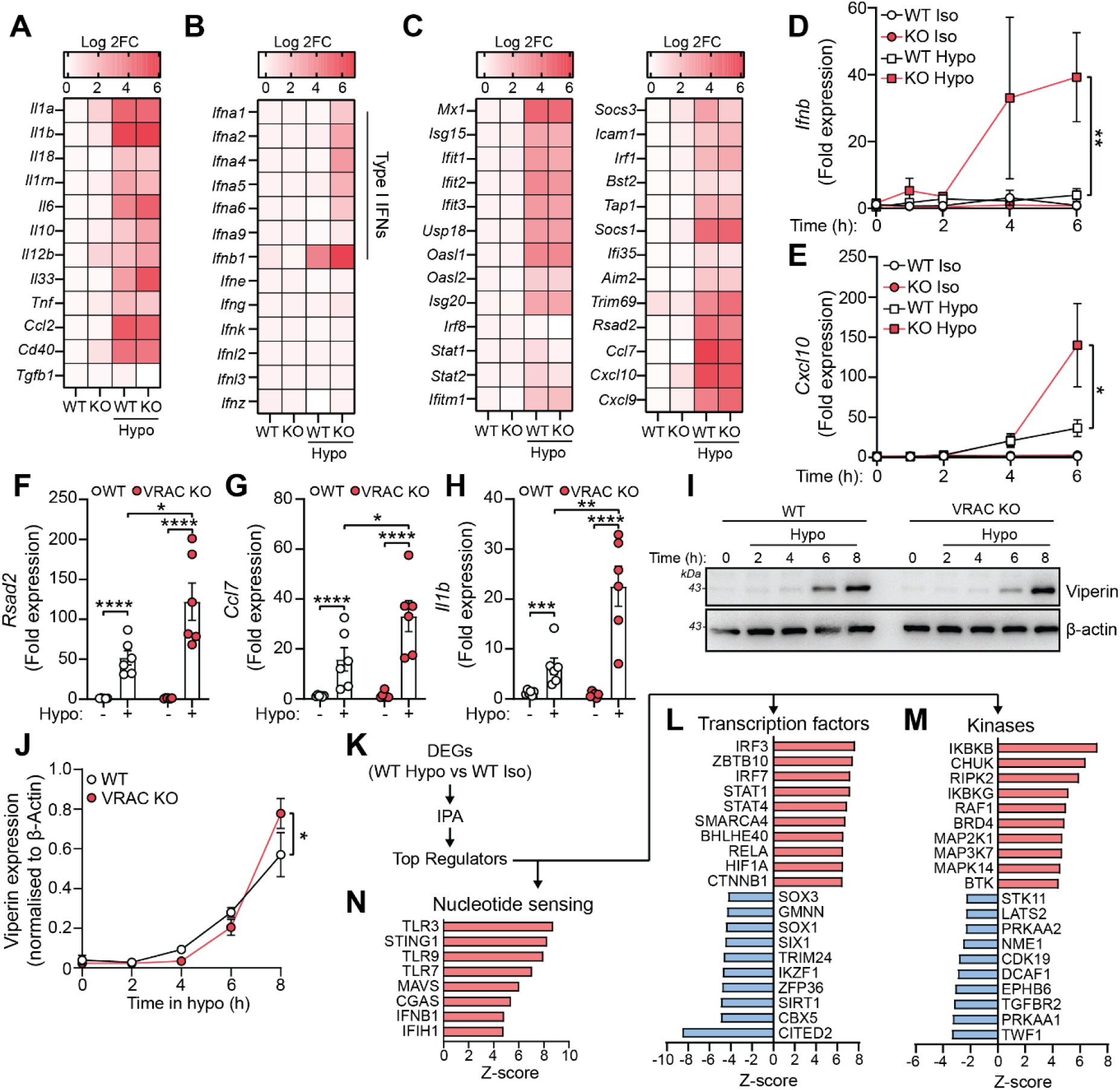
– Changes in macrophage cell volume promote inflammation through type I Interferon signalling. (**A-C**) Heatmaps of mRNA expression of cytokines (A), interferons (B) and interferon stimulated genes (C) from RNA-sequencing of wild-type (WT) or VRAC knockout (KO) BMDMs ± incubation in hypo-osmotic media (50% v/v H_2_O in DMEM). **(D-E)** qRT-PCR analysis of *Ifnb* and *Cxcl10* in WT and VRAC KO BMDMs ± incubation in hypo-osmotic media for the indicated time-points (n=4). **(F-H)** qRT-PCR analysis of *Rsad2* (F), *Ccl7* (G), and *Il1b* (H) in WT and VRAC KO BMDMs ± incubation in hypo-osmotic media for 6 hours (n=6). **(I)** Western blot for viperin in WT and VRAC KO BMDMs incubated in hypo-osmotic media for the indicated time-points (n=4). **(J)** Densitometry of (I) (n=4). **(K)** Pipeline for Ingenuity Pathway Analysis (IPA) of RNA-sequencing data (from Fig 1A) to identify potential differential upstream regulators in WT BMDMs following incubation in hypo-osmotic media. **(L-M)** Top 10 predicted upregulated and downregulated transcription factors (L) and kinases (M). **(N)** Predicted upregulated nucleotide sensing pathways. *p<0.05, **p<0.01, ***p<0.001, ****p<0.0001 determined by a one-way ANOVA (2D-E) or by a two-way ANOVA (2F-H J)) with Sidak’s post hoc analysis. Values shown are mean ± the SEM.

We then examined potential mechanisms of type I IFN signalling induction following cell volume changes. We performed ingenuity pathway analysis on our RNA-seq data from hypo-osmotic media challenged WT BMDMs to predict which kinases and transcription factors could be orchestrating this response (Fig 2K). The ten transcription factors and kinases with the highest and lowest Z-scores are shown in Figure 2L and Figure 2M. Interestingly, the regulators most likely to be activated following changes in cell volume by hypo-osmotic media were core components of antiviral signalling pathways, such as IRF3, IRF7, STAT1, and MAP3K7 (TAK1), and of pro-inflammatory signalling through NF-κB, such as RELA, HIF1A, IKBKB, IKBKG and CHUK (IKKα) (Fig 2L,M). Correspondingly, several antagonistic pathways of antiviral signalling were predicted to be downregulated, such as TRIM24. Further, there was predicted activation of sensors of cytosolic nucleotides, in particular TLR3, STING, TLR9, TLR7, MAVS, cGAS and IFIH1 which can all function upstream of IRF signalling to drive type I IFN induction (Fig 2N). These analyses suggested that changes in cell volume promotes activation of cytosolic DNA-sensing pathways leading to a type I IFN response and induction of an inflammatory cell state.

Stimulator of interferon genes (STING) is an inducer of type I IFNs^21^. Accumulation of cytosolic double stranded DNA is sensed by cyclic GMP-AMP synthase (cGAS) leading to the production of the STING agonist 2’3’cGAMP and activation of STING (Fig 3A). To test involvement of cGAS-STING in the type I IFN response to cell volume changes, we used pharmacological inhibitors of the cGAS-STING pathway in WT and VRAC KO BMDMs in response to hypo-osmotic media (Fig 3A). Matching what we saw previously, VRAC KO BMDMs, which have lost control of cell volume regulation, showed a large increase in *Ifnb* and *Cxcl10* expression following incubation in hypo-osmotic media (Fig 3B,C). Interestingly, pre-treatment with the STING inhibitor H151^22^ or the immunosuppressive oligonucleotide A151, which antagonises DNA-sensors^23^, significantly blocked the induction of *Ifnb* and *Cxcl10* in response to hypo-osmotic media (Fig 3B,C), suggesting a DNA– and STING-dependent mechanism. However, the induction of a type I IFN response was found to occur independently of cGAS, as the cGAS inhibitors Ru.251^24^ (Fig 3D,E) and 4-sulfonic calix[6]arene^25^ (Fig 3F,G) did not prevent cell swelling-induced *Ifnb* or *Cxcl10* induction. Further, antagonism of DNA-PK, another DNA-sensor proposed to function upstream of STING^26^, with NU7441 did not prevent hypo-osmotic media-induced *Ifnb* and *Cxcl10* expression (Supplementary Fig 2A,B). WT and VRAC KO BMDMs released similar levels of IFNβ when transfected with synthetic dsDNA (poly dA:dT) or treated with the STING activator 10-carboxymethyl-9-acridanone (CMA)^27^ (Fig 3H), suggesting loss of VRAC in conditions with normal cell volume regulation did not impact STING signalling. Treatment with H151, A151 or Ru.251, and to a lesser extent Nu7741, was also sufficient to block poly dA:dT-induced IFNβ release (Supplementary Fig 2C). Together, these data suggest cell volume changes drive a cGAS-independent STING signalling pathway to produce an antiviral response which is exacerbated when RVD is blocked.

**Figure 3.**
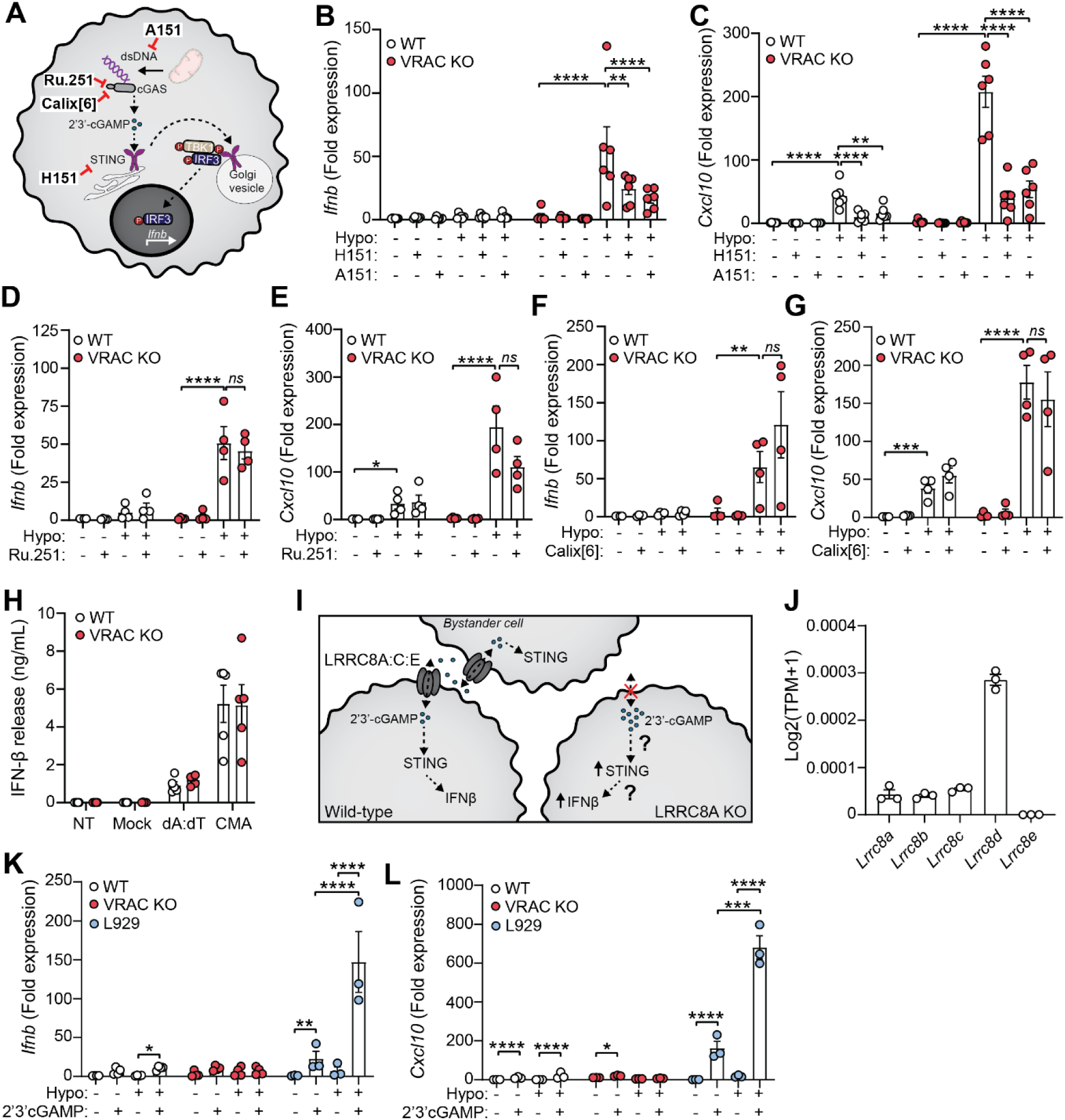
– Changes in cell volume drive IFNβ responses through a DNA-dependent STING pathway. (**A**) Schematic of the cGAS-STING pathway and inhibitors used. A151 inhibits dsDNA sensing, Ru.251 and 4-sulfonic calix[6]arene (Calix[6] are inhibitors of cGAS, and H151 inhibits STING. **(B-C)** qRT-PCR analysis of *Ifnb* (B) and *Cxcl10* (C) in WT and VRAC KO BMDMs incubated in hypo-osmotic media (50% H_2_O v/v in DMEM) for 6 hours in the presence of H151 (10 µM), A151 (1 µM) or vehicle control (DMSO 0.5% v/v) (n=4). **(D-E)** qRT-PCR analysis of *Ifnb* (D) and *Cxcl10* (E) in WT and VRAC KO BMDMs incubated in hypo-osmotic media for 6 hours in the presence of Ru.251 (10 µM) or vehicle control (DMSO 0.5% v/v) (n=4). **(F-G)** qRT-PCR analysis of *Ifnb* (F) and *Cxcl10* (G) in WT and VRAC KO BMDMs incubated in hypo-osmotic media for 6 hours in the presence of 4-sulfonic calix[6]arene (30 µM) or vehicle control (DMSO 0.5% v/v) (n=4). **(H)** IFNβ release in supernatant from WT and VRAC KO BMDMs stimulated with transfected poly dA:dT (1µg/mL), mock transfected, or treated with CMA (250 µg/mL) for 6 hours (n=5). **(I)** Schematic of proposed role for VRAC in cGAMP transport. VRAC containing LRRC8A, LRRC8C and LRRC8E can act as a conduit for 2’3’cGAMP allowing transport between extracellular space. **(J)** Number of mRNA transcripts of LRRC8 subunit genes in WT BMDMs from RNA-sequencing in Fig 1A (TPM = transcript per million). **(K-L)** qRT-PCR analysis of *Ifnb* (K) and *Cxcl10* (L) in WT and VRAC KO BMDMs, and L929 mouse fibroblasts, following incubation in isotonic or hypotonic media with or without extracellular cGAMP (5 µg/mL) for 3 hours (n=4 for BMDMs, n=3 for L929s). *p<0.05, **p<0.01, ***p<0.001, ****p<0.0001 determined by a two-way ANOVA with Dunnett’s post hoc analysis. Values shown are mean ± the SEM.

### Cell volume regulates STING function independently of cGAMP transfer

VRAC has previously been reported to modify antiviral responses by acting as a channel for 2’3’cGAMP transport across plasma membranes, facilitating immune responses by propagating STING activation into bystander cells^12–14^. Whilst we had found cGAS inhibitors were unable to prevent the type I IFN response in hypo-osmotic conditions, because of the established role for VRAC in 2’3’cGAMP transport, we explored the possibility that the increased type I IFN response that occurred in VRAC KO macrophages occurred due to defects in outward cGAMP transport, resulting in cGAMP accumulation (Fig 3I). VRAC are hexameric channels composed of a mixture of LRRC8 subunits, consisting of LRRC8A-E^7^. Whilst LRRC8A is essential for ion channel activity, and as a result RVD and volume regulation, LRRC8B-E are proposed to modify the selectivity of the pore for permeability to certain substrates^28^. In the case for optimal cGAMP transport, VRAC channels must contain LRRC8A and LRRC8E^12–14^, and high expression of LRRC8D is proposed to inhibit 2’3’cGAMP transport^13^. We therefore examined our RNA-seq dataset from our WT BMDMs (Fig 1A) to determine whether they expressed LRRC8 subunits compatible with cGAMP transport.

BMDMs expressed high levels of LRRC8D, less but equal amounts of LRRC8A, LRRC8B and LRRC8C, and interestingly did not express any transcripts for LRRC8E (Fig 3J). Therefore, the low expression of LRRC8E and high expression of LRRC8D suggests that VRAC in our BMDMs would be a poor conductor of cGAMP. To test this, we incubated WT and VRAC KO BMDMs in isotonic and hypotonic conditions for 3 hours in the presence of extracellular 2’3’cGAMP and examined induction of *Ifnb* and *Cxcl10*. Cells were incubated for only 3 hours to limit the contribution of hypo-osmotic media alone to the *Ifnb* and *Cxcl10* response. In parallel, as a positive control, we performed the same experiment in a mouse fibroblast cell line L929, which have been shown to express LRRC8E and conduct cGAMP efficiently^12^. L929 cells upregulated both *Ifnb* and *Cxcl10* in response to extracellular cGAMP and this was significantly enhanced by incubation in hypo-osmotic media (Fig 3K,L). However, WT and VRAC KO BMDMs exhibited only weak responses to extracellular cGAMP which was not enhanced by incubation in hypo-osmotic media (Fig 3K,L). Therefore, these data suggest that the induction and enhancement of STING signalling following loss of cell volume control was occurring independently of VRAC-mediated cGAMP transfer.

### Cell volume regulation modifies the inflammatory response to pathogen associated signals

Since we had established changes in cell volume promoted an inflammatory phenotype, we considered whether cell volume changes might synergise with other inflammatory triggers, such as PAMPs, to modify the inflammatory response further. This is particularly relevant as infection modifies the tissue microenvironment with signals known to modify cell volume such as oedema, hypoxia, oxidative stress, and metabolic dysfunction^10,29,30^. We therefore stimulated WT and VRAC KO BMDMs with hypo-osmotic media in the presence of the toll like receptor (TLR) agonists: Poly I:C (TLR3), LPS (TLR4), Imiquimod (TLR7) or CpG-DNA (TLR9), which drive the induction of pro-inflammatory cytokines through activation of the transcription factors NF-κB and interferon regulatory factors (IRFs)^31^. In the supernatant of stimulated BMDMs, we measured release of IL-6, a cytokine dependent on NF-κB signalling, and IFNβ, a cytokine dependent on IRF signalling to see if response to the signalling pathways had been modified. Interestingly, we found that across all PAMPs tested, IFNβ release was significantly increased by inducing changes in cell volume (Fig 4A,D). In response to poly I:C or LPS, whilst there was no difference in IFNβ release between WT and VRAC KO macrophages in isotonic conditions, stimulation in hypo-osmotic media enhanced IFNβ release in both WT and VRAC KO BMDMs, which was greater in VRAC KO cells (Fig 4A,B). Strikingly, both imiquimod and CpG-DNA only induced IFNβ release in VRAC KO BMDMs under hypotonic conditions (Fig 4C,D), suggesting loss of cell volume regulation completely altered the TLR7 and TLR9 responses. No change in IL-6 release was observed following PAMP stimulation in hypotonic conditions, except for poly I:C, which exhibited the same pattern as the Poly I:C-induced IFNβ response (Fig 4E,H). Together, these data suggest that cell volume regulation controls the inflammatory response to various PAMPs, with cell volume increases driving enhanced type I IFN responses.

**Figure 4.**
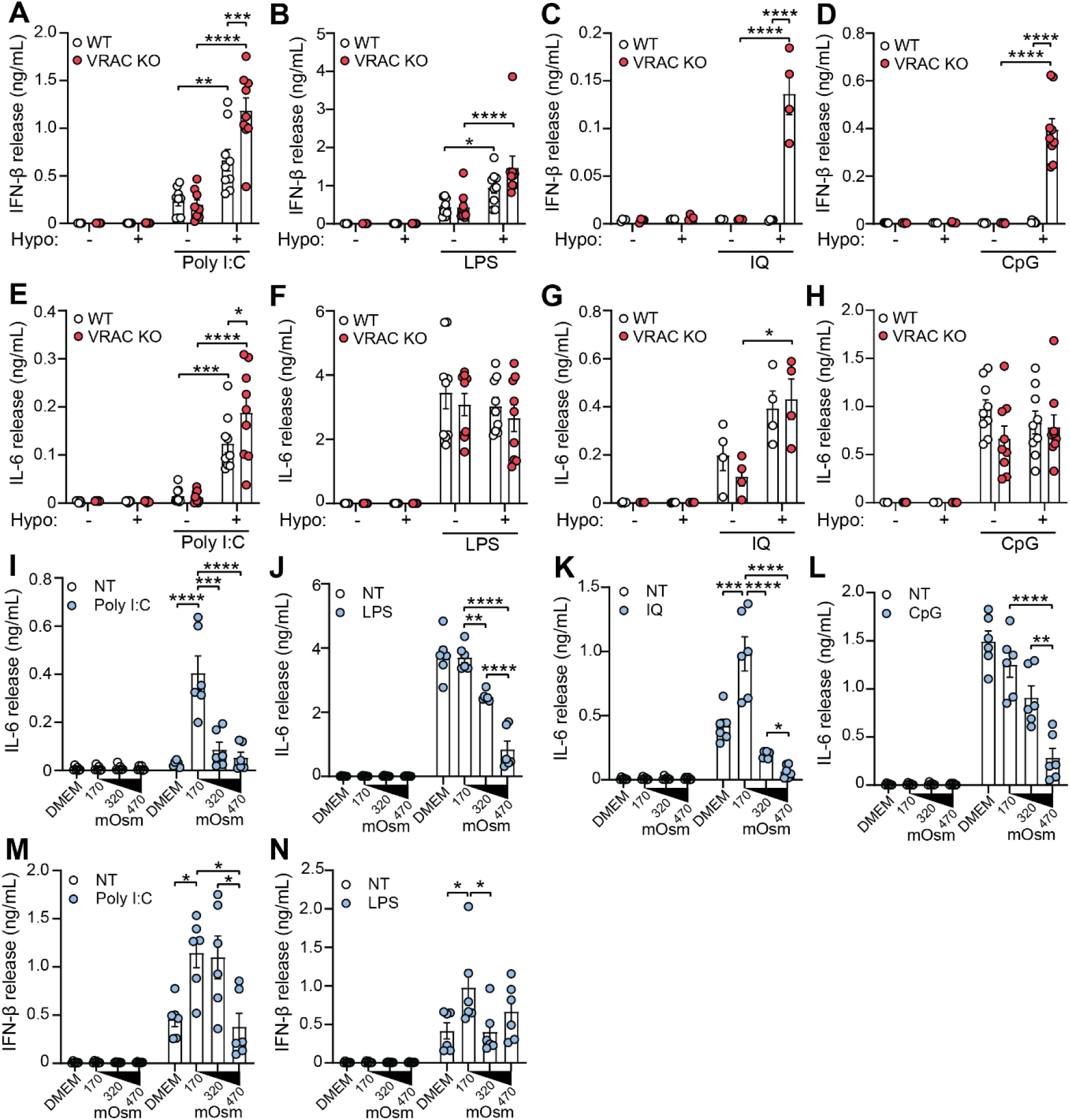
– Cell volume regulates type I interferon production in response to diverse pathogen mediated molecular patterns. (**A-D**) IFNβ release in the supernatant of WT and VRAC KO BMDMs incubated in DMEM or hypo-osmotic media (50% v/v H_2_O in DMEM) and treated with poly I:C (5 µg/mL) (A), LPS (1 µg/mL) (B), imiquimod (IQ, 25 µM) (C), or CpG-DNA (CpG, 5 µg/mL) (D) for 6 hours (n=9 for Poly I:C, LPS, CpG; n=4 for IQ). **(E-H)** IL-6 release in the supernatant of WT and VRAC KO BMDMs incubated in DMEM or hypo-osmotic media (50% v/v H_2_O in DMEM) and treated with poly I:C (5 µg/mL) (E), LPS (1 µg/mL) (F), imiquimod (25 µM) (G), or CpG-DNA (5 µg/mL) (H) for 6 hours (n=9 for Poly I:C, LPS, CpG; n=4 for IQ). **(I-L)** IL-6 release in the supernatant of WT BMDMs incubated in either DMEM, hypo-osmotic media (50% v/v H_2_O in DMEM), isotonic media (50% v/v 150 mM NaCl) or hyper-osmotic media (50% v/v 300 mM NaCl) and treated with poly I:C (5 µg/mL) (I), LPS (1 µg/mL) (J), imiquimod (IQ, 25 µM) (K), or CpG-DNA (CpG, 5 µg/mL) (L) for 6 hours (n=6). **(M-N)** IFNβ release in the supernatant of WT BMDMs incubated in either DMEM, hypo-osmotic media (50% v/v H_2_O in DMEM), isotonic media (50% v/v 150 mM NaCl) or hyper-osmotic media (50% v/v 300 mM NaCl) and treated with poly I:C (5 µg/mL) (M) or LPS (1 µg/mL) (N) for 6 hours (n=6). *p<0.05, **p<0.01, ***p<0.001, ****p<0.0001 determined by a two-way ANOVA with Sidak’s post hoc analysis 4A-H) or by a one-way ANOVA with Tukey’s post hoc analysis (4I-M). Values shown are mean ± the SEM.

Since hypotonicity-induced cell swelling and RVD modified the macrophage response to PAMPs, we investigated whether inducing a reduction in cell volume with hypertonic solutions also altered the inflammatory response. To achieve this, we incubated WT BMDMs in either DMEM, hypo-osmotic DMEM (50% v/v H_2_O), isotonic DMEM (50% v/v 150mM NaCl) or hyper-osmotic DMEM (50% v/v 300 mM NaCl), stimulated with poly I:C, LPS, imiquimod or CpG-DNA, and measured IL-6 and IFNβ release. We found that in contrast to hypotonicity, hypertonicity was effective at suppressing IL-6 release in response to poly I:C, LPS, imiquimod and CpG-DNA (Fig 4I-L). However, IFNβ release was only altered with poly I:C treatment (Fig 4M), with LPS-induced IFNβ unchanged (Fig 4N) and no IFNβ detected following imiquimod or CpG-DNA treatment (data not shown). Therefore, together these data suggest that cell volume can act to fine-tune the nature of inflammation in response to pathogens, with cell swelling promoting IFNβ signalling and cell shrinkage supressing IL-6 responses.

### Cell volume regulates inflammation in vivo

We next examined if regulation of macrophage cell volume played a role in shaping inflammatory responses *in vivo*. To do this, we used an established mouse model of macrophage activation syndrome (MAS), where a systemic inflammatory response is induced by five intraperitoneal injections of CpG-DNA over ten days^32^. This model recapitulates several features of hyperinflammatory disease, including cytokine storm and multi-organ dysfunction, with inflammation occurring throughout the body^32,33^. We therefore used the model CpG-induced MAS with our conditional VRAC KO mice (CX3CR1-cre^+/-^; LRRC8A^flox/flox^ (referred to here as VRAC^Cx3cr1-KO^) and littermate WT mice (CX3CR1-cre^-/-^; LRRC8A^flox/flox^), treated as shown in Fig 5A. Matching previous reports, repeated administration of CpG-DNA induced transient weight loss in both WT and VRAC^Cx3cr1-KO^ mice, although this was only significant in VRAC^Cx3cr1-KO^ mice (Fig 5B-C). CpG-DNA treatment also induced splenomegaly (Fig 5D), hepatomegaly (Fig 5E), and hyperferritinaemia (Fig 5F) which was the same between WT and VRAC^Cx3cr1-KO^ mice. These data suggest that loss of macrophage cell volume control did not affect the development of MAS. We then examined if the nature of the cytokine storm was altered between WT and VRAC^Cx3cr1-KO^ mice. Repeated administration of CpG-DNA caused the induction of multiple plasma cytokines, including IFNγ (Fig 5G, IL-18 (Fig 5H), TNF (Fig 5I), IL-6 (Fig 5J) and IL-10 (Fig 5K). Notably in VRAC^Cx3cr1-KO^ mice CpG treatment significantly elevated plasma IL-18, which was doubled compared to WT littermates (Fig 5H). Whilst type I IFNs were not detected in the plasma of CpG-treated mice (data not shown), IL-18 is known to be induced by interferon signalling, thus suggesting that a loss of cell volume regulation in VRAC^Cx3cr1-KO^ mice contributes to dysregulated inflammatory responses. Therefore, whilst VRAC^Cx3cr1-KO^ mice did not exhibit any significant worsening of disease phenotypes, we propose that control of macrophage cell volume does play a role in co-ordinating inflammatory responses as loss of macrophage cell volume control resulted in enhanced cytokine production.

**Figure 5.**
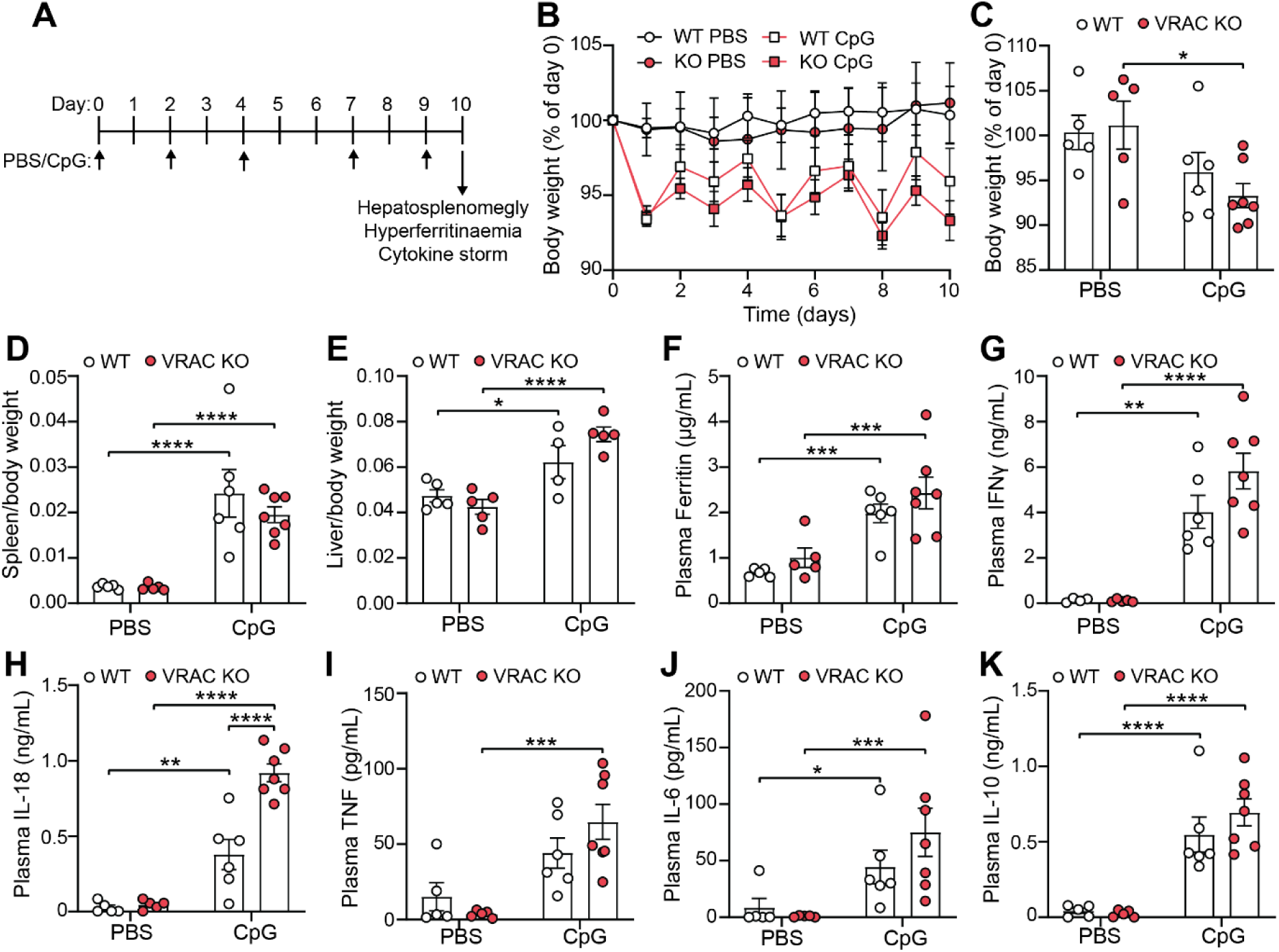
– Cell volume regulates inflammation in a murine model of hyperinflammation. (**A**) Schematic of murine model of CpG-induced hyperinflammation. CpG-DNA (2 mg kg^−1^), or PBS, was administered by intraperitoneal injection on day 0, 2, 4, 7 and 9. On day 10, tissues were taken and hyperinflammatory disease was assessed by hepatosplenomegly, hyperferritinaemia and cytokine storm. **(B)** Body weight of WT and VRAC^Cx3cr1-KO^ mice (VRAC KO) treated with PBS or CpG-DNA expressed as a percentage of day 0 (n=5-7). **(C)** Body weight at day 10 from (B) (n=5-7). **(D)** Splenic weight normalised to body weight (n=5-7). **(E)** Liver weight normalised to body weight ((n=4-5). **(F)** Plasma levels of ferritin (n=5-7). **(G-K)** Plasma concentration of IFNγ (G), IL-18 (H), TNF (I), IL-6 (J) and IL-10 (K) (n=5-7). *p<0.05, **p<0.01, ***p<0.001, ****p<0.0001 determined by a two-way ANOVA with Sidak’s post hoc analysis. Values shown are mean ± the SEM.

## Discussion

Tiered inflammatory responses related to inflammasome activation in response to increasing DAMP-mediated stress are proposed previously for mechanisms of IL-1β release^34^, and more recently for DNA sensing^35^. Related to this last example, here we propose a tiered inflammatory response to cell volume changes in macrophages. In response to extreme hypotonicity and cell swelling macrophages respond with activation of the NLRP3 inflammasome via a process that is dependent on the RVD and the Cl-channel VRAC^9,11,36^. Milder hypotonic stresses, as presented here, that are insufficient to activate NLRP3, initiate type I IFN expression and signalling, which is enhanced following loss of cell volume control. Studying hypotonicity we were also able to establish that changes in cell volume synergise with other PAMP stimuli to modify and shape inflammatory responses – highlighting cell volume regulation as a critical aspect of inflammatory signalling. Understanding disruptions in the tissue microenvironment leading to alterations in cell volume is therefore an important consideration in our understanding of disease pathogenesis.

In addition to our studies in macrophages establishing the role for VRAC in RVD-dependent NLRP3 activation^9^, VRAC-dependent RVD is also reported to be important in T-cells where loss of RVD impairs TCR signalling, resulting in impaired T-cell activation, cytokine production, proliferation, and impaired antiviral immunity to acute lymphocytic choriomeningitis virus (LCMV) Armstrong infection^37^. In addition to controlling changes in cell volume, VRAC is demonstrated to have additional important roles in antiviral immunity through acting as a plasma membrane channel for transport of 2’3’cGAMP^12–14^. Global knockout of LRRC8E, the subunit important for 2’3’cGAMP transport, inhibited induction of antiviral genes and viral clearance in mice infected with the herpes simplex virus (HSV)-1^12^. Knockout of LRRC8C in T-cells also prevented 2’3’cGAMP transport but enhanced antiviral immunity to influenza infection^38^. Furthermore, it has recently been demonstrated that several viruses, including HSV-1, VACV, and ZIKV, induce degradation of LRRC8A to prevent cGAMP transport and restrain antiviral defence^39^. The impact of virus-induced VRAC depletion on cell volume control however remains untested. Until now no studies have reported a role of LRRC8A or VRAC in macrophages regulating the RVD that contributes to IFN signalling. Here we report that VRAC influenced antiviral signalling through cell volume control in macrophages, independent of transfer of 2’3’cGAMP between cells, but dependent upon DNA sensing and STING activation, positioning cell volume control as important regulator of antiviral inflammation. Since viral infection reduces VRAC expression^39^, and consequently potentially cell volume control, it is possible that the immune system has evolved to sense a loss of cell volume control as a marker of viral infection, explaining the pro-inflammatory and antiviral response observed in this study.

Mechano-immunity has been coined to describe the interplay between cellular mechanotransduction and inflammation and is becoming increasingly recognised as an important factor in inflammatory disease^40^. Changes in the biophysical properties of the tissue environment in inflammatory conditions are known to alter the mechanical cues sensed by immune cells, resulting in altered cellular responses including cell activation, cytokine production and proliferation amongst others^41^. Pro-inflammatory signalling is known to be initiated in macrophages by mechanosensitive ion channels, such as Piezo1 and TRPV4^42,43^, and can synergise with PAMP/DAMP mediated inflammation to sensitise and augment inflammatory responses^44,45^. Here, we identified that changes in macrophage cell volume, particularly following loss of cell volume control, led to the induction of intracellular inflammatory signalling, including the NF-κB and type I IFN pathways, and augmented TLR-mediated inflammatory responses. Thus, in addition to Piezo1 and TRPV4, VRAC can also be considered a key mediator of mechano-immunity in macrophages, co-ordinating cell volume changes to translate disturbances in the inflammatory environment to tune the immune response. It is possible that the biomechanical changes that occur at the plasma membrane during the process of cell swelling and regulated volume decrease could activate additional mechanosensory pathways. Indeed, plasma membrane perturbations induced by osmotic stress can activate both Piezo1^46^ and TRPV4^47^. Interestingly, plasma membrane disruption from fusion of the viral capsid is sufficient to drive a Ca^2+-^– and STING-dependent type I IFN response^48,49^, suggesting that cell volume changes may function through similar conserved pathways. Future studies will further elucidate the interplay between VRAC and cell volume changes in the mechano-immunity response of macrophages.

In our experiments we observe that the VRAC-dependent RVD is important for NLRP3 activation in response to severe hypo-osmotic stress^9^, and that in response to milder hypoosmotic stress VRAC-dependent RVD appears to restrain inflammatory responses, highlighting a complex regulation of inflammation by cell volume change. Therapeutically, hyperosmotic solutions have been utilised to treat several conditions, predominantly following acute brain injury to reduce cerebral oedema^50–55^), but also as part of treatment for cystic fibrosis^56^, asthma^57^, bronchiectasis^58^, COVID-19^59^, and for resuscitation of critically ill patients^60^. Further, treatment with hyperosmotic solutions has been shown to limit inflammatory disease in pre-clinical disease models, including kainite-induced brain injury^36^, sepsis^15^, intracerebral haemorrhage^16^, pulmonary injury due to ischaemia-reperfusion^17^, and acute respiratory distress syndrome (ARDS)^18^, suggesting that manipulation of cell volume regulation could be harnessed to target disease processes. This is reflected in our study where hyper-osmotic solutions were capable of impairing PAMP-induced IL-6 release, as well as limiting cell swelling-induced type I IFN production and inflammasome activation. Future studies will reveal the potential for regulating VRAC-dependent RVD and cell volume changes in macrophages in disease.

## Methods

### Animals and cell culture

Conditional CX3CR1-driven macrophage specific VRAC knock out mice (*Cx3cr1^Cre/+^-;Lrrc8a^flx/flx^*) and wild-type littermates (*Cx3cr1^+/+^; Lrrc8a^flx/flx^)* were generated and maintained at the University of Manchester as previously described^9,19^). Animals were housed in ventilated cages with temperature and humidity maintained between 20–24°C and 45 to 65%, respectively, with a 12 h light-dark cycle. All procedures were performed with appropriate personal and project licenses in place, in accordance with the Home Office (Animals) Scientific Procedures Act (1986), approved by the Home Office and the local Animal Ethical Review Group, University of Manchester, and reported according to the ARRIVE guidelines. Primary bone marrow-derived macrophages (BMDMs) were isolated as previously described^25^. In brief, bone marrow was isolated from femurs and tibias, erythrocytes lysed, and cultured in DMEM (10% v/v FBS, 100 U mL^−1^ penicillin, 100 μg mL^−1^ streptomycin) supplemented with L929-conditioned media (30% v/v) for 6-7 days. BMDM cultures were fed with extra media (containing 30% v/v L929-conditioned media) on day 3. Before experiments, BMDMs were scraped and seeded overnight at a density of 1 x 10^6^ mL^−1^ in DMEM (10% FBS, 100 U mL^−1^ streptomycin/penicillin). Mouse L929 fibroblasts were cultured in DMEM (10% v/v FBS, 100 U mL^−1^ penicillin, 100 µg mL^−1^ streptomycin). L929 cells were disassociated with trypsin and seeded overnight at a density of 1 x 10^6^ mL^−1^.

### Cell stimulation

For experiments where cell volume changes were induced by hypo-osmotic shock, BMDMs were incubated in iso-osmotic DMEM (50% v/v serum free DMEM, 50% v/v sterile PBS (no Mg^2+^/Ca^2+^)) or hypo-osmotic DMEM (50% v/v serum free DMEM, 50% v/v sterile, low endotoxin, cell culture grade, H_2_O). Therefore hypo-osmotic DMEM had an osmolarity of ∼170 mOsm and iso-osmotic DMEM had an osmolarity of ∼310 mOsm.

For experiments inhibiting the cGAS-STING pathway, BMDMs were treated with either a vehicle control (DMSO, 0.5% v/v), H151 (10 µM), A151 (1 µM), Ru.251 (10 µM), 4-sulfonic calix[6]arene (30 µM) or NU7441 (100 nM) for 15 minutes prior to the addition of PBS (50% v/v) or H_2_O (50% v/v) also containing inhibitors. For experiments with chemical activation of cGAS-STING, BMDMs were incubated in serum free DMEM and stimulated with 10-carboxymethyl-9-acridanone (CMA, 250 µg mL^−1^) or lipofectamine 3000-mediated transfection of poly dA:dT (1 µg mL^−1^) for 6 hours. For experiments testing 2’3’cGAMP transport, BMDMs and L929 fibroblasts were incubated in iso-osmotic or hypo-osmotic DMEM and incubated with extracellular 2’3’cGAMP (5 µg mL^−1^) for 3 hours.

For experiments investigating TLR responses, BMDMs were incubated in either serum free DMEM, or DMEM with osmolarity altered as followed: hypo-osmotic DMEM (DMEM 50% v/v H_2_O; ∼170 mOsm), iso-osmotic DMEM (DMEM 50% v/v 150 mM NaCl; ∼320 mOsm), hyper-osmotic DMEM (DMEM 50% v/v 300 mM NaCl; ∼470 mOsm). In parallel, cells were stimulated by the addition of either poly I:C (5 µg mL^−1^), LPS (1 µg mL^−1^), imiquimod (25 µM), or CpG-DNA ODN-1826 (5 µg mL^−1^) for 6 hours.

### RNA-sequencing

RNA was isolated using a Purelink RNA miniprep kit following the manufacturer’s instructions. Total isolated RNA was submitted to the genomic technologies core facility (GTCF) at the University of Manchester. RNA samples were assessed for quality and integrity using a 2200 TapeStation (Agilent Technologies) and then libraries were generated using the TruSeq® Stranded mRNA assay (Illumina, Inc.) according to the manufacturer’s protocol. Briefly, polyadenylated mRNA was purified using poly-T, oligo-attached, magnetic beads, fragmented using divalent cations under elevated temperature and then reverse transcribed into first strand cDNA using random primers. Second strand cDNA was then synthesized using DNA Polymerase I and Rnase H. Following a single “A” base addition, adapters were ligated to the cDNA fragments, and the products then purified and enriched by PCR to create the final cDNA library. Adapter indices were used to multiplex libraries, which were pooled prior to cluster generation using a cBot instrument. The loaded flow-cell was then paired-end sequenced (76 + 76 cycles, plus indices) on an Illumina HiSeq4000 instrument. Finally, the output data was demultiplexed (allowing one mismatch) and BCL-to-Fastq conversion performed using Illumina’s bcl2fastq software, version 2.20.0.422. It is important to note that, in the RNA-Seq dataset, *Lrrc8a* RNA was still detected in KO macrophages, likely since only exon 3 of the *Lrrc8a* gene was deleted. Our previous study^19^ revealed using exon-specific primers intact expression of exons 1–2, but complete loss of exon 3 transcripts, in VRAC KO cells from *Lrrc8a^fl/fl^ Cx3cr1^cre/+^*cells.

For analysis, unmapped paired-end sequences were tested by FastQC (https://www.bioinformatics.babraham.ac.uk/projects/fastqc/). Sequence adapters were removed and reads were quality trimmed using Trimmomatic_0.36^61^. The reads were mapped against the reference mouse genome (mm10/GRCm38) and counts per gene were calculated using annotation from GENCODE M21 (https://www.gencodegenes.org/) using STAR_2.5.3a^62^. Normalization, Principal Components Analysis, and differential expression was calculated with DESeq2_1.28.1^63^. Differentially expressed genes between conditions were defined using thresholds of Log2 fold-change >0.5, adjusted p-value <.01.

### qRT-PCR

RNA was isolated using a Purelink RNA miniprep kit following the manufacturer’s instructions. cDNA was generated from isolated RNA using SuperScript III first-strand synthesis kits. qRT-PCR was performed using 5 ng cDNA, 200 nM forward and reverse primers and SYBR green mastermix according to the manufacturer’s instructions. Primer sequences used were: *Ifnb* FWD: TGGGAGATGTCCTCAACTGC, REV: CCAGGCGTAGCTGTTGTACT; *Cxcl10* FWD: CCACGTGTTGAGATCATTGCC, REV: TCACTCCAGTTAAGGAGCCC; *Rsad2* FWD: TGGTTCAAGGACTATGGGGAGT, REV: CTTGACCACGGCCAATCAGA; *Ccl7* FWD: CCCTGGGAAGCTGTTATCTTCAA, REV: CTCGACCCACTTCTGATGGG; *Il1b* FWD: CCACAGACCTTCCAGGAGAATG, REV: GTGCAGTTCAGTGATCGTACAGG; GAPDH FWD: CAGTGCCAGCCTCGTCC, REV: CAATCTCCACTTTGCCACTGC. All samples were loaded in triplicate and assayed on a 7900HT Fast Real-Time PCR machine (Applied Biosystems) for 40 cycles with standard settings. For each primer pair, a standard curve consisting of four 10-fold dilutions of neat cDNA was run in parallel to determine amplification efficiency. Sample-wise abundances were normalised to GAPDH to correct for loading differences, then further normalised to wild-type isotonic treated cells to give fold change values.

### Western blotting

BMDM lysates were generated by lysis in lysis buffer (50 mM Tris/HCl, 150 mM NaCl, 1% v/v Triton x100, pH 7.3) containing protease inhibitor cocktail. Lysates were separated by Tris-glycine SDS PAGE and transferred onto nitrocellulose or PVDF membranes using a semi-dry Trans-Blot Turbo system. Membranes were blocked in 5% w/v milk in PBS, 1% v/v Tween 20 (PBS-T) before incubated with anti-viperin antibodies (clone MaP.VIP) in 5% w/v BSA in PBS-T, overnight at 4°C. Membranes were washed three times before incubation with appropriate HRP-tagged secondary antibodies (1 h, RT), washed three times with PBS-T and then visualised with Amersham ECL prime detection reagent (GE healthcare). Chemiluminescence was visualised using a G:Box Chemi XX6 (Syngene). Membranes were re-probed for β-actin using HRP-conjugated β-actin antibodies (clone AC-15).

### *In vivo* hyperinflammation model

A mouse model of CpG-induced macrophage activation syndrome (MAS) was performed as previously described^32,33^. Conditional CX3CR1-driven macrophage specific VRAC knock out mice (*Cx3cr1^Cre/+^;Lrrc8a^flx/flx^*) and wild-type littermates (*Cx3cr1^+/+^; Lrrc8a^flx/flx^)* were used, aged 16 weeks and of both sexes. To induce MAS, mice received five intraperitoneal (I.P.) injections of CpG ODN 1826 oligodeoxynucleotides (5’-T*C*C*A*T*G*A*C*G*T*T*C*C*T*G*A*C*G*T*T-3’, where * indicates a phosphorothioate modification) (2 mg kg^−1^) or vehicle control (sterile phosphate buffered saline (PBS), 10 µL g^−1^) over ten days. Mice were randomly assigned to treatment (WT PBS n=5, KO PBS n=5, WT CpG n=6, KO CpG n=7) and researchers were blinded to both genotype and treatment for the duration of the experiment and analyses. Mice received one I.P. injection of CpG or PBS on days 0, 2, 4, 7, and 9, with tissue collected on day 10, 24 hours after the last injection. Animals were weighed daily. Mice were anaesthetised, blood was collected via cardiac puncture and plasma isolated by spinning at 1500*xg* for 15 minutes at 4°C, before storage at –70°C. Following cardiac puncture, animals were transcardially perfused with PBS before collection and weighing of the spleen and liver. Plasma ferritin levels were assessed via ELISA (Abcam, ab157713), plasma IL-18 levels were assessed via ELISA (Invitrogen, BMS618-3), and plasma IFNγ, TNF, IL-6 and IL-10 were determined by a LEGENDplex multiplex assay (mouse inflammation panel, Biolegend, 740446), all according to the manufacturer’s instructions.

### Regulatory volume decrease (RVD) assay

5 x 10^4^ BMDM were seeded out into black-walled 96-well plates overnight. Cells were loaded with 10 μM calcein-AM (10 µM, 1 hr, 37°C), washed three times with DMEM and then rested for 30 min to allow calcein to equilibrate. After three further washes with serum-free DMEM, cells were imaged on an Eclipse Ti microscope (Nikon) equipped with a stage incubator maintaining 37°C and 5% CO_2_. Calcein fluorescence was captured in the GFP channel using low laser power to minimize phototoxicity and bleaching. Live imaging was conducted using point-visiting to image all conditions simultaneously. One image was captured every 2 min, with hypotonic shock induced at 5 min by adding endo-toxin free distilled H_2_O (50% v/v) for a final osmolarity of ∼170 mOsm kg^−1^. For quantification, fluorescent images were processed in Fiji, using a rolling-ball background subtraction (50 pixels ball size) to remove non-cell associated fluorescence. The average pixel intensity of each frame was then measured and normalized to the value of the first frame, yielding F/F_0_ curves. For statistical testing, AUC values of the curves were calculated in Prism.

### Quantification and statistical analysis

Data are presented as mean values plus the SEM. Accepted levels of significance were *p < 0.05, **p < 0.01, ***p < 0.001, ****p < 0.0001. Statistical analyses were carried out using GraphPad Prism (version 8). Equal variance and normality were assessed with Levene’s test and the Shapiro–Wilk test, respectively, and appropriate transformations were applied. Groups containing normally distributed data were analysed using either a one-way ANOVA with Sidak’s or Tukey’s post hoc analysis, or a two-way ANOVA with Sidak’s, Tukey’s or Dunnett’s post hoc analysis as appropriate. n represents experiments performed on individual animals or cells acquired from individual animals.

