## Supplementary figures for "Cell volume tunes macrophage innate inflammatory responses through promoting type I interferon signalling"

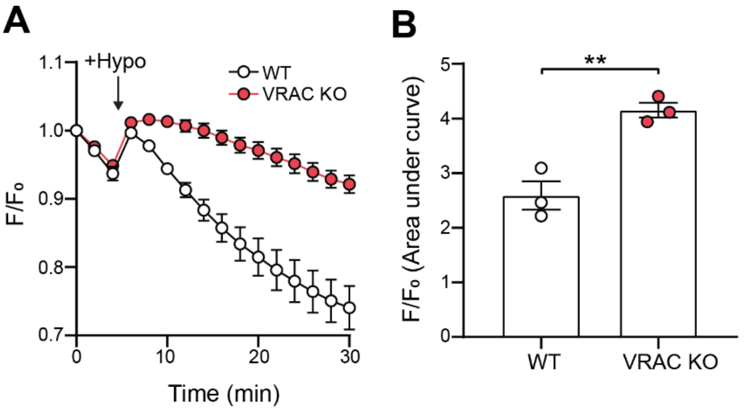


**Supplementary Figure 1 – Related to Figure 1: Knockout of LRRC8A removes regulatory volume decrease in BMDMs. (A)** Regulatory volume decrease measured by calcein fluorescence in WT or VRAC knockout (KO) BMDMs incubated in hypo-osmotic media (50% v/v $H_2O$ in DMEM, ~170 mOsm $kg^{-1}$), added at the indicated time point (n=3). These data were obtained from the same cultures of BMDMs used for RNA-sequencing analysis in figure 1. **(B)** Area under the curve analysis of (A). **p<0.01 determined by an unpaired t-test. Values shown are mean ± the SEM.

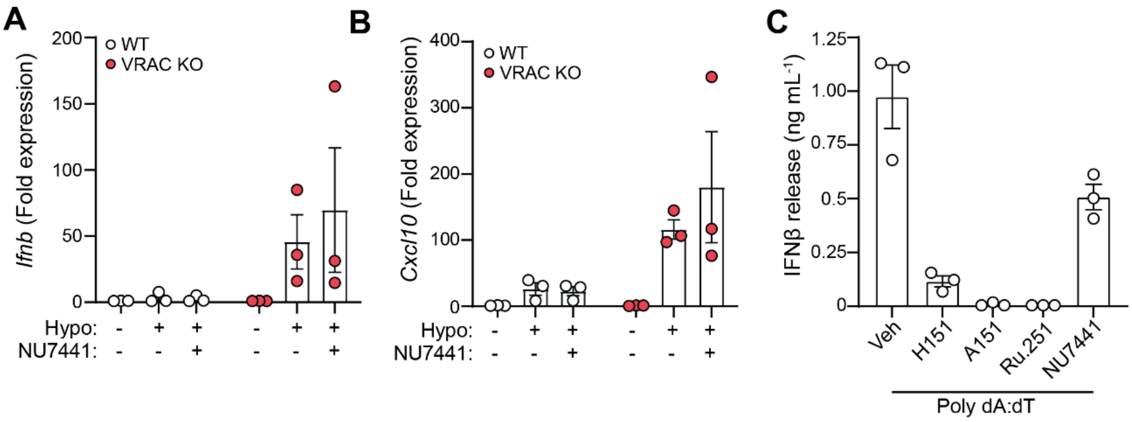


**Supplementary Figure 2 – Related to Figure 3: Changes in cell volume drive IFNβ responses through a DNA-dependent STING pathway. (A-B)** qRT-PCR analysis of *Ifnb* (A) and *Cxcl10* (B) in WT and VRAC KO BMDMs incubated in hypo-osmotic media (50% $H_2O$ v/v in DMEM) for 6 hours in the presence of Nu7441 (100 nM), or vehicle control (DMSO 0.5% v/v) (n=3). **(C)** IFNβ release in supernatant from WT BMDMs stimulated with transfected poly dA:dT (1µg/mL) in the presence of H151 (10 µM), A151 (1 µM), Ru.251 (10 µM), NU7441 (100 nM) or vehicle control (DMSO, 0.5% v/v) for 6 hours (n=3). Values shown are mean ± the SEM.
